# Synchronous seasonality in the gut microbiota of wild wood mouse populations

**DOI:** 10.1101/2021.10.15.464528

**Authors:** K.J. Marsh, A.M. Raulo, M. Brouard, T. Troitsky, H. M. English, B. Allen, R. Raval, J.P. Webster, S. C. L. Knowles

**Affiliations:** Department of Pathobiology and Population Sciences, The Royal Veterinary College, University of London, Hawkshead Campus, Hatfield, Hertfordshire, AL9 7TA; Department of Zoology, University of Oxford, Mansfield Road, Oxford, Oxfordshire, OX1 3SZ; Department of Life Sciences, Imperial College London, Silwood Park, Buckhurst Road, Ascot, SL5 7PY; College of Life and Environmental Sciences, University of Exeter, Penryn Campus, Cornwall, UK

**Author notes:** Author for correspondence: Kirsty J Marsh, College of Life and Environmental Sciences, University of Exeter, Penryn Campus, Cornwall, UK, TR10 9FE.

**Keywords:** Bacteroidales, core, individuality, lab vs wild, microbiome, mouse, seasonality, 16S

## Abstract

1. The gut microbiome performs many important functions in mammalian hosts, with community composition shaping its functional role. However, what factors drive individual microbiota variation in wild animals and to what extent these are predictable or idiosyncratic across populations remains poorly understood.
2. Here, we use a multi-population dataset from a common rodent species (the wood mouse, *Apodemus sylvaticus*), to test whether a consistent set of ‘core’ gut microbes is identifiable in this species, and to what extent the predictors of microbiota variation are consistent across populations.
3. Between 2014 and 2018 we used capture-mark-recapture and 16S rRNA profiling to intensively monitor two wild UK mouse populations and their gut microbiota, as well as characterising the microbiota from a laboratory-housed colony of the same species.
4. Although broadly similar at high taxonomic levels and despite being only 50km apart, the two wild populations did not share a single bacterial amplicon sequence variant (ASV). Meanwhile, the laboratory-housed colony shared many ASVs with one of the wild populations from which it is thought to have been founded decades ago. Despite strong taxonomic divergence in the microbiota, the factors predicting compositional variation in each wild population were remarkably similar. We identified a strong and consistent pattern of seasonal microbiota restructuring that occurred at both sites, in all years, and within individual mice. While the microbiota was highly individualised, seasonal convergence in the gut microbiota among individuals occurred in late winter/early spring.
5. These findings reveal highly repeatable seasonal gut microbiota dynamics across distinct populations of this species, despite divergent taxa being involved. Providing a platform for future work to understand the drivers and functional implications of such predictable seasonal microbiome restructuring, including whether it might provide the host with adaptive seasonal phenotypic plasticity.

## Introduction

The gastrointestinal tracts of vertebrates harbour complex microbial communities known as the gut microbiota, that can perform a wide range of important functions for the host. These include regulating the immune system (Round & Mazmanian, 2009), extracting nutrients from otherwise indigestible parts of the diet (Flint et al., 2012), and defence against pathogens (Buffie & Pamer, 2013). With such important roles, one might expect these symbiotic communities to be under strong host influence, such that individuals of a given species harbour a characteristic and relatively invariant community. Yet the emerging picture from microbiome research firmly contradicts this. Vertebrate gut microbiotas are typified by immense compositional variation, both among individuals and within individuals over time. Each individual’s gut microbiota constitutes a diverse community of microbes shaped by environmental influences such as diet, habitat and xenobiotics, microbial interactions, metacommunity-level processes such as transmission among hosts, and mechanisms of host selection, arising for example through physiological differences across host genotypes or age. The relative importance of these different ecological processes in shaping wild animal microbiomes remains a key open question.

In humans, cross-population microbiome comparisons have shown that only a small set of ‘core’ gut bacteria are consistently detected across individuals within a population, and fewer are shared across populations (Falony et al., 2016; Li et al., 2013; Qin et al., 2010; Tap et al., 2009). Rather, the human gut microbiota is highly individualised, with each person harbouring a characteristic microbial fingerprint which is distinct from that of others over time (Bergström et al., 2014; De Muinck & Trosvik, 2018; Faith et al., 2013; Turnbaugh et al., 2009). However, given the unique ecology of industrialised humans, whether such patterns apply to non-human wild vertebrates remains poorly understood. Cross-population studies in the wild examining to what extent taxa are shared by conspecifics across geographical space, and at what scale this might occur, are rare (Grieneisen et al., 2019; Lankau et al., 2012; Linnenbrink et al., 2013; Smith et al., 2015; Funosas et al., 2021). Moreover, most wild animal microbiota studies are cross-sectional or group-level, with few that have characterised the microbiota of repeat-sampled individuals. Such longitudinal data are crucial for understanding microbiota stability at both the individual and population level, and how this may be shaped by fluctuations in the environment.

A growing number of studies have revealed strong seasonal dynamics in the gut microbiota of wild animals (Kobayashi et al., 2006; Williams et al., 2013; Fogel, 2015; Springer et al., 2017; Amato et al., 2014; Sun et al., 2016; Liu et al., 2019; Ren et al., 2017; Maurice et al., 2015; Orkin et al., 2019). Pronounced seasonal microbiota dynamics have also been observed among humans leading traditional, hunter-gatherer lifestyles (Davenport et al., 2014; Smits et al., 2017). These findings contrast with those from industrialised humans, where microbiota composition is considered relatively stable in adulthood (Faith et al., 2013). The causes of such seasonal dynamics are not fully understood, but existing studies have implicated seasonal shifts in diet (Amato et al., 2014a; Ren et al., 2017), or hibernation (Carey, Walters, & Knight, 2013; Sommer et al., 2016). However, in many studies the fine-scale seasonal dynamics are not well-understood as temporal resolution is relatively course (e.g. winter vs. summer), and the extent to which seasonal dynamics are idiosyncratic or repeatable across years or populations of a species remains unknown. Understanding the repeatability of such seasonal dynamics across time and space is a key step towards understanding their potential functional significance. If such temporal dynamics are general across populations and years, this would strengthen the support for a broadly applicable underlying process (for example a predictable environmental or intrinsic host seasonal change), and allow for the possibility that such changes could provide the host with repeatable adaptive seasonal plasticity (Alberdi et al., 2016; Amato et al., 2014a).

Here, we present a longitudinal analysis of gut microbiota dynamics across two wild wood mouse (*Apodemus sylvaticus*) populations, together with a comparison from a wild-derived captive colony of the same host species. Wood mice are an ideal study system for questions relating to the wild host microbiome, as they are a common species that are amenable to identifiable capture-mark-recapture protocols. Furthermore, a previous study on a UK wood mouse population has described seasonal shifts in gut microbiota composition, occurring between summer and autumn (Maurice et al., 2015). We provide comprehensive analyses of intra- and inter-individual host level variation and elucidate the factors associated with this variation in each wild population.

## Methods

### Sample collection

Rodents were live-trapped using a capture-mark-recapture study design in two mixed deciduous woodlands approximately 50km apart in the UK: Wytham Woods, Oxfordshire (‘Wytham’, 51° 46’N,1°20’W) and Silwood Park, Berkshire (‘Silwood’, 51°24’N, 0°38’W). Fieldwork methods were consistent across sites, with trapping performed for one night every 2-4 weeks across all seasons. In Silwood, trapping occurred on a single 2.47ha grid over a one-year period (Nov 2014 to Dec 2015), while in Wytham trapping took place over a three-year period (Oct 2015 to Oct 2018), initially on a 1ha grid that was then expanded to 2.4ha in October 2017. Wood mice (*Apodemus sylvaticus*) were the most commonly caught rodent at both sites, with yellow-necked mice (*Apodemus flavicollis*) and bank voles (*Myodes glareolus*) also captured. Small Sherman live traps were baited with six peanuts and a slice of apple, with sterile non-absorbent cotton wool provided for insulation. Traps were set in alternate 10×10m grid cells at dusk and collected at dawn. All newly captured rodents were injected subcutaneously with a PIT tag for identification purposes, with ear notches used as a back-up form of identification. The following information was recorded for all captures: species, ID, sex, age (juvenile, sub-adult or adult, according to size and pelage characteristics), reproductive state (active or inactive according to signs of reproduction such as testes size in males, perforation, pregnancy or lactation in females), body mass (to the nearest 0.1g), subcutaneous fat score (0-4, measured by palpating the lower spine and hips). All animals were released within the 10×10m grid cell of their capture. Faecal samples (approximately 40-300mg) were collected from the bedding material with sterilized tweezers, and frozen at -80°C within 10 hours of trap collection for molecular work. Traps that showed any sign of animal contact (both traps that held captured animals and those where an animal had interfered with but not triggered it) were washed thoroughly with 20% bleach solution between trapping sessions to prevent cross-contamination. Live-trapping work was conducted with institutional ethical approval, and at Wytham under Home Office licence PPL-I4C48848E.

Faecal samples were also collected from an outbred, captive colony of wood mice kept at the University of Edinburgh, and included for comparison to wild populations. This colony was initiated over 30 years ago at the University of Oxford (Department of Zoology) using wild-caught mice from the local area (presumed to be Wytham Woods, where department members were trapping rodents at the time, though this has not been confirmed) and bred there for several decades (Hughes et al., 2010). Captive wood mice included in this study were sampled as part of a diet shift experiment carried out in 2017 under Home Office Project license 70/8543. Further details about the colony and experiment are given in the Supporting Information.

### 16S rRNA gene sequencing

Genomic DNA was extracted from faecal samples using Zymo Quick-DNA faecal/soil Microbe 96 (96-well plate format) extraction kits, according to manufacturer instructions. A 254bp region of the bacterial 16S rRNA gene (V4 region) was amplified using primers 515F/806R (Caporaso et al., 2011; Table S1). Library preparations followed a two-step (tailed-tag) approach with dual-indexing (D’Amore et al., 2016) and sequencing was performed on an Illumina® MiSeq with 250bp paired-end reads, at the Centre for Genomic Research in Liverpool. Samples were extracted in 17 batches and sequenced in 5 lanes, with some Wytham and colony samples extracted and sequenced together, while samples from Silwood were extracted and sequenced separately with minor adjustments to the protocol (Supporting Information). To avoid further batch effects influencing results, wild samples from each trapping session and colony samples from each timepoint and animal were evenly distributed across extraction plates. Full details of 16S rRNA sequencing methods are provided in Supporting Information.

### Bioinformatics

Raw sequence data were processed through the DADA2 pipeline (v1.6) in R to infer amplicon sequence variants (ASVs; Callahan et al., 2016, 2017). This was done separately for each Miseq run, but using identical parameters. In brief, reads were trimmed and filtered for quality, ASVs inferred and putative chimeras removed before taxonomic assignment performed using the v128 SILVA reference database. Full details of the bioinformatics pipeline are in Supporting Information. A single *phyloseq* object (McMurdie & Holmes, 2013) containing data from all three populations was then created for further processing and analyses. ASVs taxonomically assigned as chloroplast or mitochondria were removed, after which the dataset contained 11,404 ASVs. The R package ‘iNEXT’ (Chao et al., 2014; Hsieh et al., 2016) was used to examine sample completeness and rarefaction curves, which showed that sample completeness plateaued at approximately 8000 reads (Fig. S1). 31 samples with read counts below this threshold were removed, leaving data from 1052 samples and 281 mice overall (Wytham: n= 448 samples from 178 individuals; Silwood: n=253 from 75 individuals; captive colony: n=351 from 28 individuals), with a mean read count of 40,478 (range 8841-150,932). For beta diversity analyses, further taxon filtering was performed by retaining only those ASVs with more than 1 copy in at least 1% of samples, to guard against the influence of potential PCR or sequencing artefacts, or contaminants. After this additional ASV filtering step, the dataset contained 2662 ASVs.

### Statistical analyses

All statistical analyses were performed in R version 3.5.3 (R Core Team, 2019).

#### Defining core taxa

To delimit core taxa, we estimated each ASV’s population-wide *prevalence* as the proportion of samples in which it was detected, in a dataset containing one randomly selected sample per individual, averaged over 100 iterations. We also estimated each ASV’s within-individual *persistence*, as the proportion of samples per repeat-sampled individual it was detected in, and taking the mean across mice sampled at least three times (n=57 in Wytham, with mean N captures 4.86±1.94 s.d., n=39 in Silwood with mean N captures=5.95±1.95 s.d.).

#### Alpha-diversity analyses

ASV richness was estimated using the R package ‘breakaway’ (Willis & Bunge, 2015), using a dataset for which taxon filtering was limited to the removal of ASVs assigned as chloroplast or mitochondria.

#### Beta-diversity analyses

ASV read counts were normalised to relative abundance per sample before calculating pairwise dissimilarity indices among samples (Bray-Curtis dissimilarity; McKnight et al., 2019), which was subsequently used in principal coordinates analysis (PCoA) and permutational analysis of variance (PERMANOVA) per population. To assess the decay of within-individual community similarity over time (time-decay), we modelled the log-linear change in community structure (pairwise Bray-Curtis dissimilarity) with the number of days between samples. Community dissimilarities were converted to similarities (1-dissimilarity) and log-transformed. The rate of change at different time lags was calculated by dividing the Bray-Curtis dissimilarity by the time between sampling points to obtain an average rate of change for each time lag (Faith et al., 2013; Shade et al., 2013). To compare inter- and intra-individual variation in gut microbiota composition, pairwise dissimilarity values were compared across different types of sample pair, using Wilcoxon tests with 1,000 permutations: those collected from the same individual at different timepoints (‘Same mouse’) and those collected from different individuals within the same trapping session (‘Same date’).

To analyse temporal change in microbiota composition, generalised additive mixed models (GAMM) were run in package ‘mgcv’, using sample scores along the first axis of a Bray-Curtis PCoA (PC1) as the response variable. A cyclic cubic spline was fitted for day of the year to model within-year seasonal patterns, with corresponding sample-level host and methodological terms included as covariates (age, sex, reproductive status, body mass, body condition, read count). Year and MiSeq run were also fitted as covariates in analyses for Wytham. Animal ID was included as a random factor to control for repeated measures. The same model structure was used to assess seasonal patterns in microbial richness.

To test the relative effects of host, environmental and methodological factors on variation in gut community structure, a marginal PERMANOVA was performed using the *adonis2* function in package ‘vegan’ (Oksanen et al., 2019) with 999 permutations, on a subset of data including one randomly chosen sample per individual, to avoid pseudoreplication. Subsequent tests for homogeneity of variance between groups of significant terms were performed using the *betadisper* function. To circumvent the decrease in sample size from subsetting to one sample per individual, we also performed a partial redundancy analysis (pRDA) on Hellinger-transformed community data, using the same explanatory terms as used in the PERMANOVA, but with individual ID used as a condition. The marginal significance of the explanatory terms were then calculated under the reduced model.

To identify bacterial taxa (ASVs) involved in detected trends, we used Random Forest models with default parameters (package ‘randomForest’). ‘IncNodePurity’ was used as a measure of ASV importance.

## Results

### Assessing core microbes across populations

Across all three (wild and captive) populations, the gut microbiota of wood mice was broadly comparable in terms of proportional abundances of high-level taxa, being dominated by the orders Lactobacillales, Clostridiales and Bacteroidales (Fig. 1a). At the order level, Wytham mice had a slightly higher relative abundance of Bacteroidales, while Silwood mice had a slightly higher relative abundance of Lactobacillales. The extent to which a set of ‘core’ microbes common to all populations was identifiable declined as higher levels of bacterial taxonomic resolution were considered (Fig. 1b). Nine phyla were detected in total, eight of which were detected in all populations; Spirochaetes was only found in Wytham and the colony while Elusimicrobia was found only in Silwood. The majority (74%) of identified bacterial orders were common to all populations and similarly 62% of identified genera were shared by all three populations, wild and lab (Fig. 1b). However, strong taxonomic divergence was detected among populations at the ASV level; no ASVs were shared between the two wild mouse populations, nor between Silwood wild mice and the captive colony (Fig. 1b). This finding was unaltered when taxon filtering was minimal and limited to the removal of reads assigned as chloroplast/mitochondria. In contrast, more than half the ASVs seen in wild mice from Wytham were also detected in the captive colony (52.1% of Wytham and 95% of Colony ASVs; Fig. 1b). Differences in the proportion of taxa shared by populations were unaltered when these comparisons were made using samples sequenced in the same vs. different runs (Fig. S2), indicating they are very unlikely to be an artifact of the fact Silwood samples were sequenced separately. On average, a pair of mice from Wytham and the colony shared 8.4% ASVs (mean Jaccard Index), while the proportion of shared ASVs within a population was slightly higher (Wytham=11.5%, Colony=23.2%, Silwood=18.0%). The ASVs common to Wytham and colony mice came from a broad range of bacterial families, representative of those observed in each population (Fig. S3).

**Figure 1.**
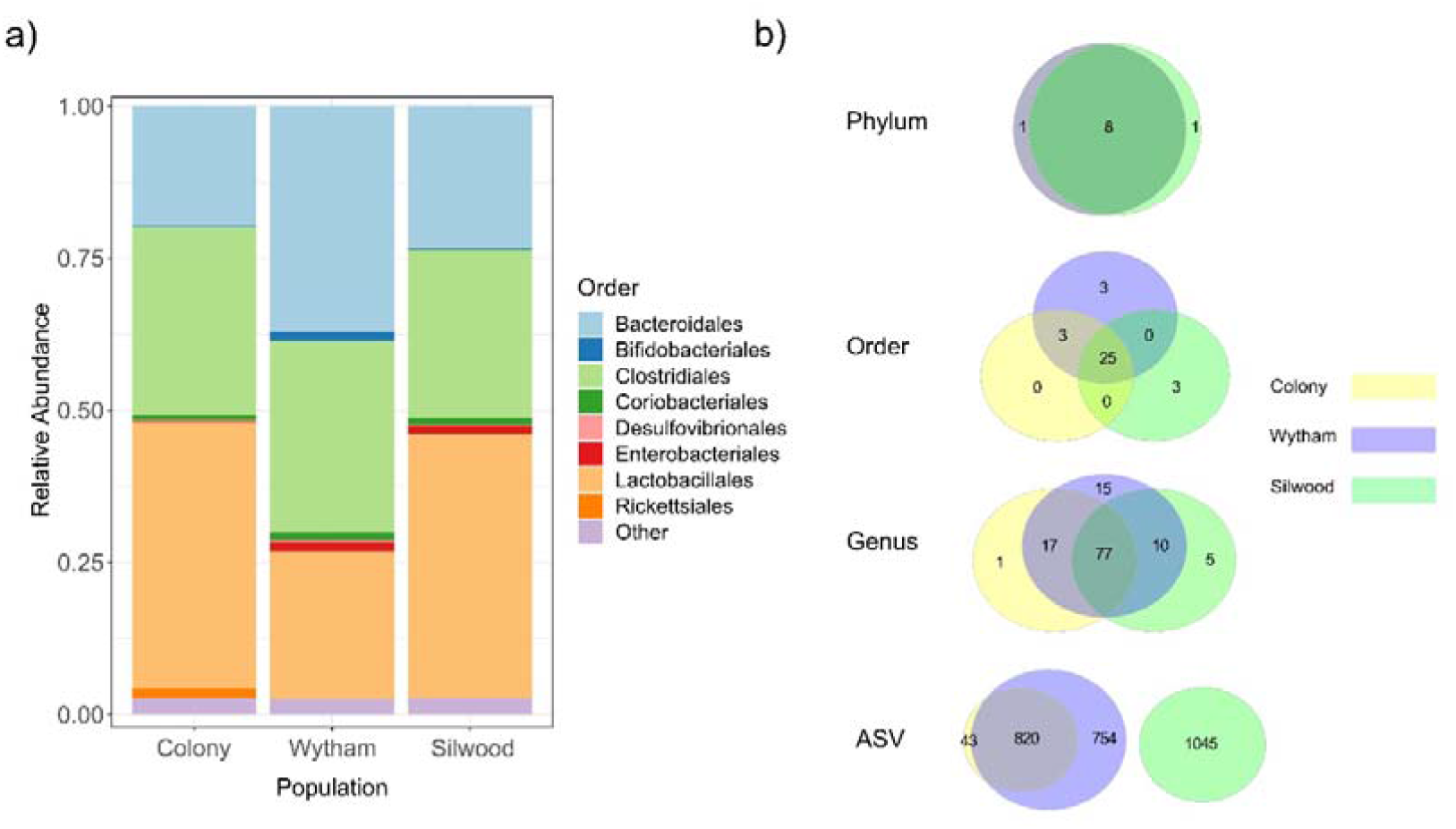
Gut microbiota composition and taxon sharing across populations of wood mice. **a)** Comparison of microbiota composition at the Order level in samples from a captive colony (n=351) and two wild populations, Wytham (n=448) and Silwood (n=253). **b)** Euler diagrams showing the number of shared and unique taxa across populations at four taxonomic levels: Phylum, Order, Genus and ASV (amplicon sequence variant). Circle size is proportional to the total number of taxonomic units, and circles are coloured by population: Colony (yellow), Wytham (blue), Silwood (green).

### Assessing core microbes within populations

There were no ASVs shared by all samples within each population, i.e. no population-specific ‘core microbiota’. To explore how we might define a set of ‘core’ microbiota members using a less strict criterion than universal colonisation, we first examined the relationship between ASV prevalence across hosts and persistence within them. In both wild populations, there was a strong positive correlation between the prevalence and persistence of ASVs (Spearman’s rank correlation; Wytham; rho=0.932, p<0.001, Silwood; rho=0.946, p<0.001, Fig. S4). ASVs that were both prevalent and persistent (>50% for each) formed a taxonomically biased subset, being enriched for the order Bacteroidales (and slightly for Lactobacillales) compared to the total set of ASVs in each wild population (Fig. S4).

Defining core taxa using both prevalence and abundance thresholds (present in at least 60% samples at 0.1% relative abundance or more) identified a similar set of ASVs in each population. In Wytham, this gave a set of 23 core ASVs, belonging to Bacteriodales S24-7 group (n=16, 69%), Peptococcaceae (n=1), Ruminococcaeae (n=3) and Lactobacillaceae (n=3). Only two of these ‘core’ ASVs could be identified to genus level and belonged to *Lactobacillus* and *Ruminiclostridium 9*, respectively. Using the same core definition, the Silwood population contained 21 core ASVs, belonging largely to the Muribaculaceae (n=11) as well as Ruminococcaceae (n=4), Lactobacillaceae (n=4), Helicobacteraceae (n=1) and Coriobacteraceae (n=1).

### Predictors of gut microbiota composition

In both wild populations, month was the strongest predictor of gut microbiota composition, explaining approximately twice as much variation as all host factors combined in marginal PERMANOVAs (Table 1). Although population age structure varies seasonally (Fig. S5), the seasonal effect detected here is independent of this, as host age was controlled for in the model (Table 1). Microbiota composition also differed between years in Wytham. Weaker effects of host factors (body mass and reproductive status) were detected, but each in only one of the two populations (Table 1). Dispersion tests indicated that the effect of year may have been influenced by dispersion differences in Wytham while the effect of month may also contain some influence of dispersion in Silwood (Table 1). Results from the pRDA on Bray-Curtis dissimilarity agreed with those from PERMANOVAs, in that month and year strongly predicted gut microbiota variation in both wild populations, while measured host factors did not (Table S3). However, the pRDA also revealed that individual ID (as a condition) explained around half the total variation in each dataset (55.2% in Wytham and 48.1% in Silwood).

**Table 1.** Predictors of microbiota composition in two populations of wild mice. Results are shown from a marginal PERMANOVA on Bray-Curtis dissimilarity values. One randomly selected sample per individual was included in the model (Wytham; n=128 and Silwood; n=75). p-values <0.05 are in bold. For significant terms (factors only), tests for multivariate homogeneity of group dispersions were carried out and the results presented alongside the PERMANOVA. The interaction between sex and reproductive status was fitted in a separate model including this interaction term, results for all other terms are from a model without this interaction.

| Variable | Wytham |  |  |  | Silwood |  |  |  |
| --- | --- | --- | --- | --- | --- | --- | --- | --- |
|  | df | F | p | Partial R<sup>2</sup> | df | F | p | Partial R<sup>2</sup> |
| Read count | 1 | 1.338 | 0.104 | 0.009 | 1 | 0.629 | 0.895 | 0.007 |
| MiSeq run | 3 | 1.460 | **0.006** | 0.030 |  |  |  |  |
| Month | 11 | 1.758 | **0.001** | 0.133 | 10 | 1.584 | **0.001**‡ | 0.187 |
| Year | 3 | 2.591 | **0.001**† | 0.053 |  |  |  |  |
| Sex: Reproductive status | 1 | 0.840 | 0.699 | 0.006 | 1 | 0.835 | 0.628 | 0.009 |
| Sex | 1 | 0.908 | 0.598 | 0.006 | 1 | 0.786 | 0.684 | 0.009 |
| Reproductive status | 1 | 1.320 | 0.126 | 0.009 | 1 | 1.731 | 0.066 | 0.020 |
| Age | 2 | 1.001 | 0.428 | 0.014 | 2 | 0.930 | 0.563 | 0.022 |
| Body mass | 1 | 1.382 | 0.065 | 0.010 | 1 | 1.018 | 0.382 | 0.012 |
| Body condition | 4 | 0.944 | 0.642 | 0.026 | 4 | 0.928 | 0.615 | 0.044 |

† and ‡ indicate terms for which dispersion tests indicated significant differences in dispersion among groups, with p=0.003 and p=0.01 respectively

### Repeatable seasonal restructuring of the microbiota

The major axis of microbiota compositional variation (PC1 from a Bray-Curtis PCoA, explaining 12.89% of variation in Wytham and 20% in Silwood) shifted strongly between July, October and February, and this seasonal pattern was remarkably consistent across both mouse populations (Fig. 2a, c; Table S2). PC1 was not significantly predicted by any other host or methodological fixed effects (Table S2). In both populations, seasonal changes in PC1 within repeat-sampled individuals typically tracked population-level seasonal shifts, with few exceptions (Fig. 2b, d; paired t-tests for Wytham Jun-Aug vs. Sept-Nov; t=-5.152, p<0.001, n=14, Sept-Nov vs. Jan-March; t=2.676, p=0.015, n=20, for Silwood Jun-Aug vs. Sept-Nov; t=-3.23, p=0.006, n=16, Sept-Nov vs. Jan-March; t=0.250, p=0.805, n=26). Of the few individuals that were not consistent with the population-level patterns, there were no unique or unusual observations in the measured host traits, though repeat captures were too few between these months to formally test factors associated with individual differences in the direction or magnitude of seasonal change in PC1. The pattern of seasonal change in microbiota composition was similar when using the PC1 of Jaccard-based PCoA (a presence/absence-based distance metric) as the response, although in Silwood the Jaccard-based PC1 explained a lower proportion of gut community variation than Bray-Curtis PC1 (Fig. S6). This shows that gut microbiota seasonality primarily consists of changes in relative abundance rather than turnover, as can be seen in Sankey plots showing the flux of ASVs between seasons (Fig. S7).

**Figure 2.**
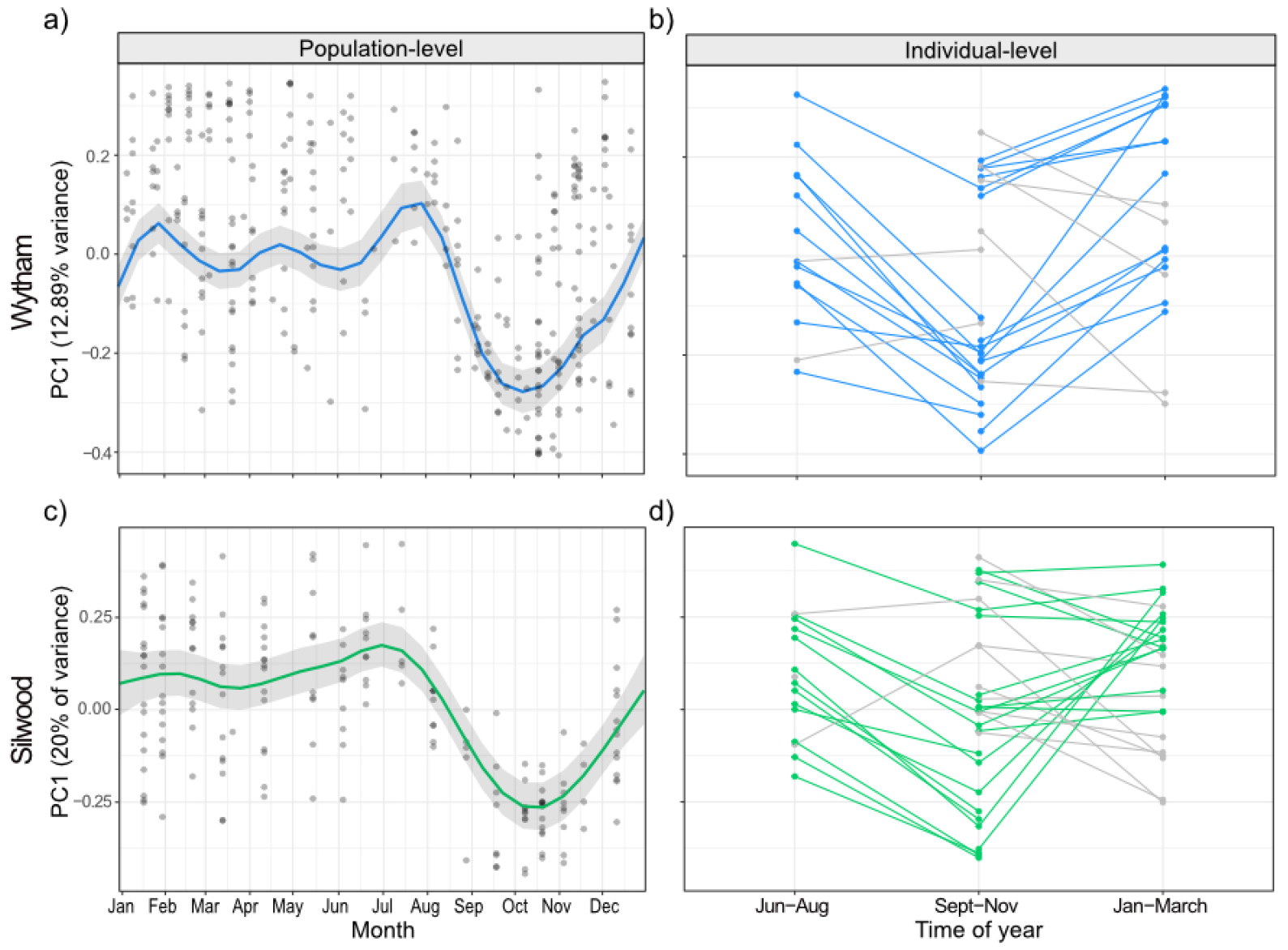
Seasonal restructuring of the wood mouse gut microbiota in two wild populations. **a)** and **c)** Seasonal dynamics in population-level PC1 value (the first axis of a principal coordinates analysis on Bray-Curtis dissimilarity values) in Wytham (top, blue) and Silwood (bottom, green). Data from Wytham mice come from a 3-year period (Oct 2015-2018), while Silwood mice were sampled for one year (November 2014-15). Predicted values and 95% confidence intervals for the smoothed day of the year from generalised additive mixed models (GAMMs) are plotted. **b)** and **d)** Changes in PC1 within repeat-sampled individual mice typically track the population-level seasonal shifts in both populations (coloured lines), with some exceptions (grey lines).

Despite this prominent seasonal shift in microbiota composition, the relative abundances of bacterial families did not change dramatically across the year (Fig. S8). Exploratory analyses also showed that while in Silwood the second and third PCoA axes showed some temporal fluctuations during the same periods that PC1 shifted (oscillation between July and December), in Wytham other PCoA axes besides PC1 did not show strong seasonal variation (Fig. S9).

Seasonal changes in mean gut microbiota richness were weaker compared to seasonal changes in composition. In models where richness was the response, day of the year was non-significant for Wytham mice, and in Silwood was significant though explained a relatively small amount of variation compared to similar analyses of composition (Fig. S10, Wytham: approximate significance of smoothed date term F=0.00, edf<0.001, p=0.516, adjusted R^2^=0.09, n=328; Silwood: approximate significance of smoothed date term; F=1.767, edf=4.722, p<0.001, adjusted R^2^=0.14, n=185).

We used random forest regressions to explore which bacterial taxa might drive the consistent seasonal changes in PC1 seen in each wild population. For both populations, the top 6 ASVs predicting PC1 values belonged to three bacterial families: Lactobacillaceae, Muribaculaceae and Ruminococcaceae, with the exact order of importance for these families differing slightly between populations (Fig. 3). In Wytham, the model explained 92.11% of variation in PC1 values, and the most important ASVs predicting PC1 belonged to Ruminococcaceae, followed by Muribaculaceae, Lactobacillaceae and Bifidobacteriaceae (Fig. 3). In Silwood, the model explained 93.31% in PC1 and all top four ASVs in predicting PC1 belonged to Lactobacillaceae, followed by ASVs belonging to Muribaculaceae and Ruminococcaceae (Fig. 3). The bacterial ASVs most strongly associated with variation along Silwood PC2 and PC3 belonged to Lactobacillaceae, Ruminococcaceae, Lachnospiraceae and Muribaculaceae (Fig S11).

**Figure 3.**
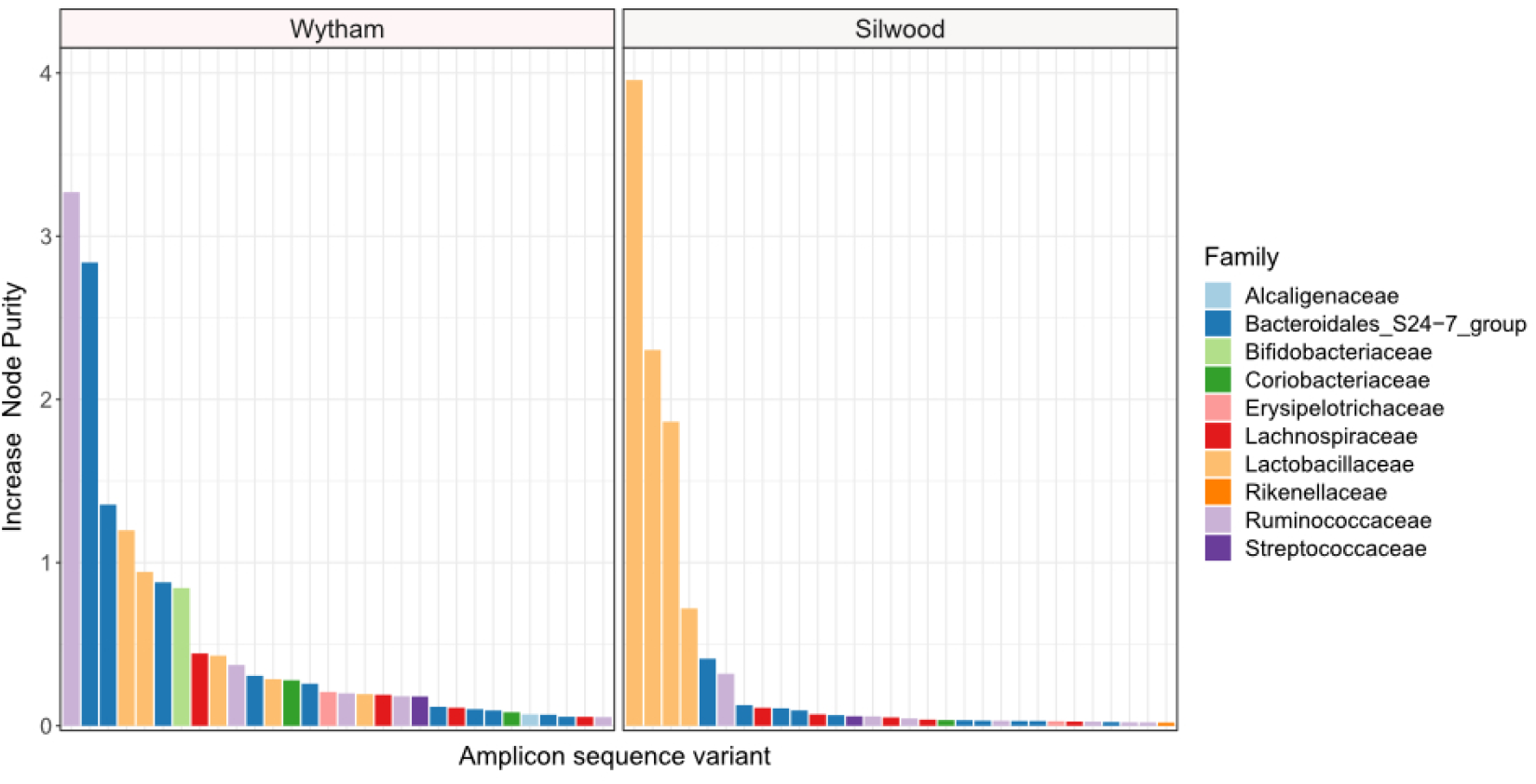
The importance of bacterial taxa in driving consistent seasonal patterns in PC1. (first axis of a Bray-Curtis PCoA) in two wild wood mouse populations. Random forest regressions (RFR) were used to identify bacterial amplicon sequence variants (ASVs) important for predicting PC1, which has a strong seasonal signal, in each population. In Wytham (left), the RFR could explain 92.11% of variation in PC1 using the relative abundance of ASVs as predictors, while the Silwood the RFR could explain 93.31% of variation in PC1. The ‘IncNodePurity’ was used as a measure of feature importance in the models, and the top 30 strongest ASVs (highest IncNodePurity) are shown, with the bacterial family each ASV in associated with indicated by colour.

### Individuality and seasonal convergence in the gut microbiota

Despite strong and repeatable seasonal shifts in population-level gut microbiota composition (Fig. 2), we nonetheless detected a large degree of individuality in the microbiota of wild mice. In each wild population, individuals were, on average, more similar in gut microbiota composition (Bray-Curtis dissimilarity) to themselves at other time-points than to other mice sampled on the same day (Fig. 4a; permutation tests; Wytham observed U-statistic=4222156, p<0.001; Silwood observed U-statistic=1802548, p<0.001). This signal of individuality decayed with increasing sampling interval (Fig. 4b; log-linear model for Wytham; F=107.6_(1,1268)_, p<0.001, adjusted R^2^=0.078, Silwood; F=17.51 _(1,918)_, p<0.001, adjusted R^2^=0.018), with the rate of decay highest at short time intervals and less strong as the sampling interval increased (Table S4).

**Figure 4.**
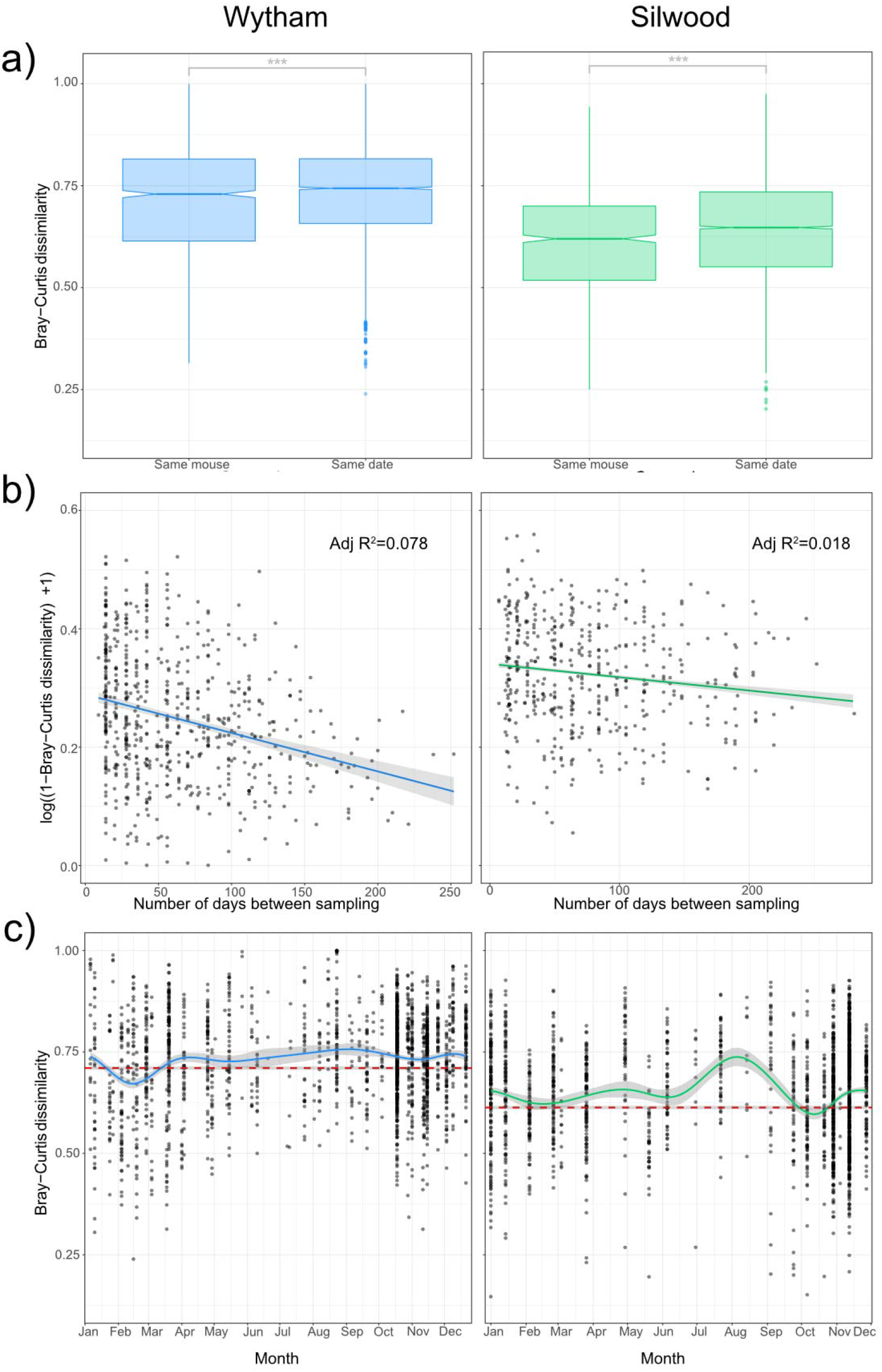
Intra- and inter-individual variation in gut community structure varies with time in two wild populations. **a)** Pairwise Bray-Curtis dissimilarity between samples from the same host at different times (Same mouse) and those taken from different hosts on the same day (Same date) were used to compare intra- and inter-individual variation in Wytham (blue) and Silwood (green) wild populations. Significant differences between groups were tested with permutational Wilcoxon tests and are denoted by asterisks (***; p<0.001, **; p<0.01, *; p<0.05, NS; p>0.05) **b)** Intra-individual variation decays with sampling interval. Pairwise community similarity (1-Bray-Curtis dissimilarity) between pairs of samples collected from the same individual host, in mice that were captured three or more times in two separate populations (Wytham; n=277 samples from 57 hosts and Silwood; n=197 samples from 39 hosts) are plotted against the number of days between the samples being collected. Community similarity is log-transformed and the relationship is fitted using a log-linear model. **c)** Inter-individual variation across the year was visualised by plotting the Bray-Curtis dissimilarity between individuals per trapping session with a loess smoothing line for each wild population. The mean (±se) intra-individual Bray-Curtis dissimilarity per population is shown as a dashed reference line in red.

Although there was a strong signal of individuality overall (intra-individual variation is on average lower than inter-individual variation), this does not account for temporal dynamics in the community composition. We therefore also assessed seasonal changes in inter-individual variation (Bray-Curtis dissimilarity) to determine whether the signal of individuality remains strong throughout the year. In general, there tended to be more convergent gut microbiotas between individuals in winter but more inter-individual variation in the summer months (Fig. 4c). This inter-individual variation remained higher than the average within-individual variation for most of the year (Fig. 4c), showing that the signal of individuality remains consistently strong. However, in Wytham, the seasonal convergence of gut microbiota composition seen in late winter/ early spring was sufficiently strong to override the signal of individuality (Fig. 4c); mice sampled in February were more similar to each other than they were to themselves at other times of the year (permutation test; observed U-statistic= 223084, p<0.001). A weaker convergence was observed in the Silwood population at this same time of year (late winter/ early spring), which nullified the signal of individuality i.e. mice caught at this time were no more similar to themselves across the year than they were to others sampled simultaneously (permutation test; observed U-statistic= 33916, p= 0.712). This reduction in inter-individual variation was observed again in October in Silwood (permutation test; observed U-statistic= 30240, p= 0.046) but not in Wytham (Fig. 4c).

## Discussion

Here we have provided an in-depth examination of natural variation in the gut microbiota of a wild mammalian host, the wood mouse, using detailed longitudinal analyses from multiple populations and years. We found no universal core microbial community (shared by all members of the species) when considering bacterial ASVs, genera or families across all populations. The proportion of bacterial taxa shared between host populations decreased with increasing taxonomic resolution, and at the finest resolution of ASVs there was a high degree of population-specificity in the taxa identified, suggesting a high amount of turnover in the species/strains present even between populations 50km apart. Geographic variation in gut microbial communities has been observed in a number of host species, with processes such as differences in diet and/or host genetics (Goertz et al., 2019; Smith et al., 2015; Suzuki et al., 2019), environmental differences such as soil properties (Grieneisen et al., 2019) or ecological drift (Lankau et al., 2012; Linnenbrink et al., 2013; Stothart et al., 2020) cited as drivers of this variation. In contrast, the captive colony of wood mice retained approximately half of the diversity of the wild population from which it likely originated, indicating a slow rate of taxon loss over time in captivity with minimal input of new microbes as has been observed in other mouse systems (Kohl & Dearing, 2014; Sonnenburg et al., 2016). The patterns observed here were not simply an artefact of the sequencing methodology, as the subsets of Wytham and colony samples sequenced in the same or different runs shared similar proportions of taxa in common. The degree of shared ASVs between the wild and captive populations studied here could be driven either by the degree of host or microbial genetic divergence between populations (highest between the two wild populations, but lower between the colony and Wytham), or by variability of environmental microbial input, which is high between wild populations but limited in captivity.

Associations between microbiota structure and host body mass (Sommer et al., 2016; Turnbaugh et al., 2006), age (Bennett et al., 2016; Degnan et al., 2012) and reproductive status (Amato et al., 2014b) have been detected previously. However, here we show that sampling month remains a strong predictor of microbiota variation after accounting for these intrinsic host factors, and seasonality is therefore likely driven by external factors in this system. These seasonal microbiota changes were much stronger than any host-associated effects examined, with sampling month explaining more than twice the variation in community structure than any other variable tested, and more than all host-related variables combined. This suggests that associations with host factors are relatively weak compared to external drivers of gut microbiota variation, in line with studies on other wild animal populations (Goertz et al., 2019; Ren et al., 2017) and humans (Rothschild et al., 2018).

By sampling with high temporal resolution throughout multiple years, we detected a strong seasonal oscillation in microbiota structure that consistently occurred between July and February. This seasonal shift occurred independently of host age, sex or reproductive status, and was also observed within individuals. The timing and form of this seasonal microbiota restructuring was remarkably consistent across both populations and all years examined, and strongly resembled patterns detected in a two-year study of a separate UK wood mouse population (Maurice et al., 2015). Other studies in wild animals have similarly reported a dominant role of season in restructuring the gut microbiota compared to host factors (Kobayashi et al., 2006; Williams et al., 2013; Fogel, 2015; Springer et al., 2017; Amato et al., 2014a; Sun et al., 2016; Liu et al., 2019; Ren et al., 2017; Maurice et al., 2015). However, this is the first time such seasonality has been shown to have such a consistent pattern across multiple populations and years. The striking repeatability of this seasonal pattern across populations suggests common drivers are at play. These could include predictable seasonal shifts in diet, parasitic infection, host physiology or social behaviour (David et al., 2014; Perofsky et al., 2017; Kreisinger et al., 2015). Hibernation has been linked to restructuring of the gut microbiota in ground squirrels and brown bears (Carey et al., 2013; Sommer et al., 2016). Although wood mice do not hibernate, they can enter short periods of torpor during colder months, with decreased body temperature and metabolic activity. However, this explanation of seasonal restructuring in the wood mouse gut microbiota is unlikely, as the major seasonal restructuring begins in summer. Therefore, seasonal shifts in diet are a strong candidate mechanism. Wood mice are omnivorous species that in woodland habitats rely on nuts which mature in summer and are cached for consumption until the following spring, such that their diet shows a strong increase in this material during the autumn/winter (Watts 1968). This predictable influx of nuts to the diet could be what drives consistent seasonal restructuring across populations and years in woodland habitats, a hypothesis that could be tested in this system through direct measurement of seasonal diet-microbiota links in future.

Despite strong seasonal changes in the gut microbiota that were observed consistently across individuals and populations, a high degree of individuality in the gut microbiota was still detectable. On average, individuals were more similar to themselves over time than to other individuals sampled simultaneously. Such individuality could be explained by genetic differences (Benson et al., 2010), or persistent environmental or behavioural differences, for example in habitat, dietary preferences or parasite burden. This individuality signal weakened as sampling interval increased, suggesting that temporal changes (e.g. environmental or aging-related) can affect the strength of this individual gut microbial signature (Faith et al., 2013). We also found seasonal variation in the degree of microbiota similarity between hosts, with the gut microbiota being more similar among mice caught in winter and early spring. This convergence of gut microbiotas in February-March was sufficiently strong to reduce the signal of individuality and even temporarily override it within the Wytham population. This could be due to a more homogenous diet at this time of year, when wood mice are thought to eat mainly cached seeds and nuts, compared to later spring/summer, when a more diverse array of both plant and animal items is available (Watts 1968). Alternatively, seasonal convergence could be driven by population density and/or social interactions, which vary seasonally and could influence rates of microbial transmission between hosts (Raulo et al., 2021; Wolton, 1985). Seasonal convergence between individual hosts in relation to the signal of individuality in the gut microbiota has not been previously examined. However, this could have consequences for the host population in terms of how well the gut microbiota can provide resilience to perturbations such as altered food availability or pathogen invasion at different times of the year.

The repeatability of seasonal microbiota dynamics across populations observed here is particularly striking given these two host populations possessed no bacterial ASVs in common. Some, but not all, of the higher order taxa associated with seasonal changes were consistent across populations. For instance, Lactobacillaceae ASVs were implicated in both Wytham and Silwood, consistent with previous work (Maurice et al., 2015), while members of Ruminococcaceae, Bacteroidales S24-7 group, Bifidobacteriaceae and Lachnospiraceae were important in Wytham only. Thus across populations, different bacterial taxa appear to respond synchronously to the same seasonal change. This suggests there may be a level of functional redundancy in the gut microbial taxa of wood mice that respond seasonally at broad geographical scales. Further functional studies, for example using metagenomic or metatranscriptomic approaches, would be valuable to illuminate what seasonal functions these microbes might perform for the host, and the potential significance of such microbiome shifts in providing hosts living in variable environments with adaptive seasonal plasticity.

## Acknowledgements

This work was funded by a NERC fellowship (NE.L011867/1) to SCLK and a Royal Veterinary College (RVC) studentship to KJM (supervised by SCLK and JPW). Our thanks go to Tim Coulson for enabling the fieldwork at Wytham Woods, and to Amy Pedersen for enabling the sample collection in the captive wood mouse colony.

## Author Contributions

This work was supported by a NERC fellowship (NE.L011867/1) to SCLK and funding from the European Research Council (ERC) under the European Union’s Horizon 2020 research and innovation programme (grant agreement n° 851550). KJM was supported by a Royal Veterinary College (RVC) studentship (supervised by SCLK and JPW). Our thanks go to Tim Coulson for enabling the fieldwork at Wytham Woods, and to Amy Pedersen for enabling sample collection in the captive wood mouse colony.

## Data Availability Statement

All relevant data will be made available on a suitable repository.

## Supporting Information

Supp_Mat_file1.docx

