## Supplemental_Information_1 for "Synchronous seasonality in the gut microbiota of wild wood mouse populations"

**Supplementary Methods**

*Sampling of captive mice*

Captive wood mice were sampled from an outbred laboratory colony of wood mice, kept under semi-barrier conditions at the University of Edinburgh, Institute of Evolutionary Biology. Colony wood mice are kept in individually-ventilated cages (IVCs) under specific pathogen-free conditions, and fed *ad libitum* on Certified Rodent Diet 5002 (LabDiet). The sampled colony mice included here were involved in a diet shift experiment, in which they were singly housed for 4 weeks and fed one of three diets: standard sterile lab chow, or chow supplemented with irradiated peanuts, dried mealworms, or a mix of both. Faecal samples were collected from individuals at 14 regularly spaced time-points throughout the diet shift experiment, and frozen at -80°C within 7 hours of collection.

*16S rRNA gene sequencing*

Genomic DNA was extracted from 1179 wood mouse faecal samples using Zymo Quick-DNA faecal/soil Microbe 96 kits, according to manufacturer instructions. Colony samples were extracted across 7 plates along with the first two years of samples collected from Wytham, the last year of samples from Wytham were extracted across a further 6 plates along with samples from a separate study (unpublished data) and the Silwood samples were extracted across a further 4 plates. Samples from each trapping session were spread across extraction plates for Wytham and Silwood, and colony samples from each timepoint and animal were split across extraction batches. The lysis stage was optimised for maximum DNA yield with the Qiagen Tissuelyser II set at a frequency of 25/s for 25 minutes. The lysis stage for the 4 Silwood plates was conducted using a tube format bead beater in batches of 24 samples with a frequency of 3000rpm for 8 minutes. 80 µl of DNA was eluted and stored at 4°C while tests of extraction success were being carried out, and subsequently at -80°C for long-term storage. One negative (water) extraction control and one mock community extraction (ZymoBIOMICS™ Microbial Community Standard) were included in each plate of 96 extractions, for quality control purposes. Extracted DNA concentration (ng/µl) was determined using the HS dsDNA kit on a Qubit® 2.0 fluorometer. Amplicon generation and library preparation followed a two-step approach with dual-indexing (D’Amore et al. 2016) and was carried out in conjunction with the Centre for Genomic Research (CGR), University of Liverpool. The first step involved use of Illumina-adapted bacterial primers targeting the V4 region of the bacterial 16S rRNA gene (Table S1; Caporaso et al., 2011), and the second stage involved addition of Illumina barcodes to enable multiplexing samples in each sequencing run. Test PCRs were performed using 35 cycles of the first round PCR, to assess amplification success of the samples and potential contamination. Test PCRs were performed in 10µl containing 5 µl KAPA mix, 0.125µl each of forward and reverse primers at 10uM, 2.25µl PCR water and 2.5µl neat DNA extraction (or negative/positive control). The PCR conditions were as follows; 98°C for 2 minutes, 35 cycles of 95°C for 20s, 65°C for 15s and 70°C for 30s, followed by 72°C for 5 minutes. PCR products were visualised on 2% agarose gel stained with GelRed, and no amplification was observed in negative controls. Based on the results of this test PCR, DNA extractions were diluted either 0- or 4- fold such that after 35 cycles a mid-strength band would be expected. DNA extractions were then subjected to two replicates of the first round PCR as above except with 20 cycles. These PCR products were pooled by sample and submitted to CGR for 15 cycles of the second round PCR (to add barcodes), amplicon size selection, pooling and sequencing. Samples were sequenced on Illumina® MiSeq runs (384 samples per run) with paired end 2x250bp sequencing. Five Miseq runs were performed; runs 1 and 2 contained samples from Wytham and the colony, runs 3 and 4 contained Wytham samples and run 5 contained Silwood samples.

**Table S1. Primer sequences for amplification of the V4 16S rRNA region.** Round 1 primers (Caporaso et al., 2011) are in red, and the recognition sequence to allow the second round PCR is in blue. Round 2 Illumina adapter sequences are italicised and example barcodes (Illumina indices) are in bold.

| **PCR round** | **Primer name** | **Primer sequence** |
| --- | --- | --- |
| 1 | 515F | 5' ACACTCTTTCCCTACACGACGCTCTTCCGATCTNNNNNGTGCCAGCMGCCGCGGTAA 3' |
| 1 | 806R | 5' GTGACTGGAGTTCAGACGTGTGCTCTTCCGATCTGGACTACHVGGGTWTCTAAT 3' |
| 2 | N501F (example barcode) | 5' *AATGATACGGCGACCACCGAGATCTACAC***TAGATCGC**ACACTCTTTCCCTACACGACGCTC 3' |
| 2 | N701R (example barcode) | 5' *CAAGCAGAAGACGGCATACGAGAT***TCGCCTTA**GTGACTGGAGTTCAGACGTGTGCTC 3' |

*Bioinformatics using the DADA2 pipeline*

Raw sequence reads were processed through the DADA2 v1.6 pipeline in R following the online tutorial (<https://benjjneb.github.io/dada2/tutorial_1_4.html)>. First, sequence quality was examined across the lengths of the reads to inform trimming parameters. Five samples from each sequencing run were inspected in order to exclude sequence regions after which sequence quality declined; forward reads were truncated at 200bp and reverse reads at 175bp using the *filterandTrim* function. In the same step, primer sequences were removed from the 5’ end of the reads using trimLeft=24 to remove the first 24bps (this was chosen in an initial analysis using cutadapt (Martin 2011) which showed that trimming at this length would remove the primer sequence from the vast majority of reads), otherwise the standard filtering parameters were used. Between 48-98% of reads per sample survived trimming and filtering. After dereplication and inference of amplicon sequence variants (ASVs) using the DADA2 algorithm trained on estimated forward and reverse read error rates (Callahan et al. 2016), paired reads were merged and sequences of extreme length were filtered out, retaining only those with approximately the expected length (between 246-251bp), before removal of putative chimeras. Overall, an average 95% of reads per sample survived the pipeline across all the MiSeq runs. Taxonomy was assigned to the ASVs using the v128 SILVA reference database (formatted for DADA2; silva_nr_v128_train_set.fa.gz).

**Figure S1. Sample completeness and rarefaction curves**. Curves are shown for each sample after taxonomic filtering (removal of sequences assigned to chloroplast and mitochondria) and prior to abundance filtering (removal of sequences of low prevalence/abundance) generated in the R package iNEXT.

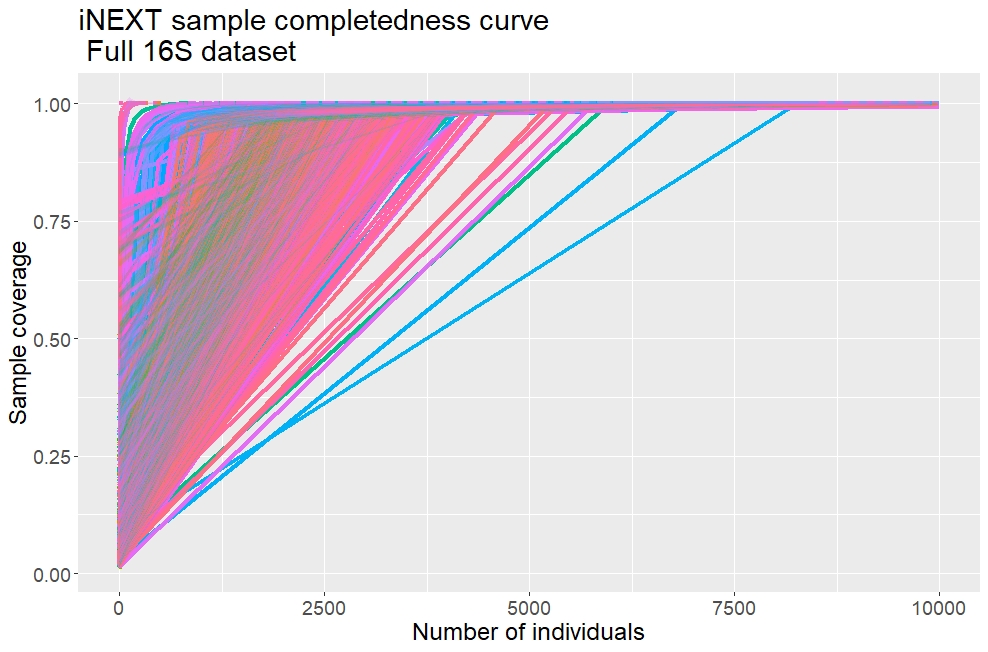

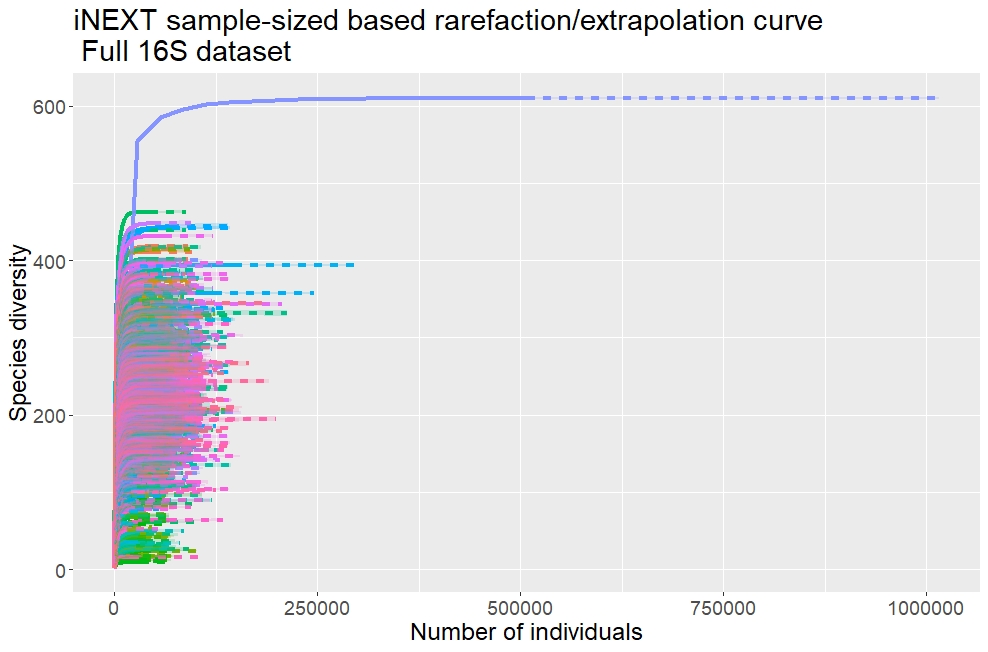

**Supplementary Results**

**Figure S2. Extent of gut microbe sharing across wild and colony mice does not depend on sequencing run.** The percentage of microbial taxa shared between populations at different taxonomic levels. Samples from a wild population, Wytham (W, n=448) and a captive colony (C, n=351) were compared. (tog)=samples that were sequenced in the same MiSeq run (sep)=samples that were sequenced in separate runs.

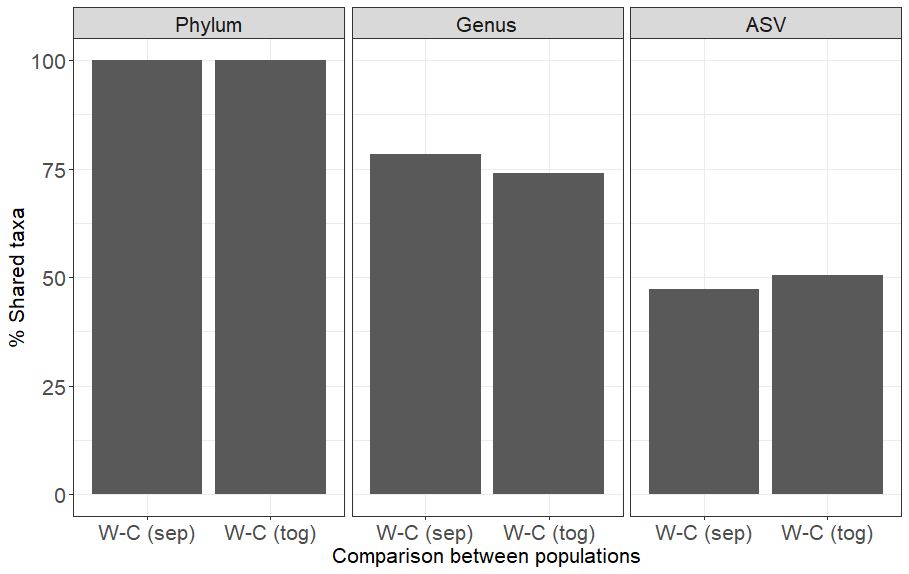

**Figure S3. The taxonomic distribution of bacterial ASVs common or unique to mice from Wytham Woods or the captive colony.** The taxonomic distribution of bacterial amplicon sequence variants (ASVs) that were present in each population, common to both, or unique to one or the other are compared. The proportion of ASVs in each subset assigned to different bacterial families are shown, with the total number of ASVs indicated at the top of each bar.

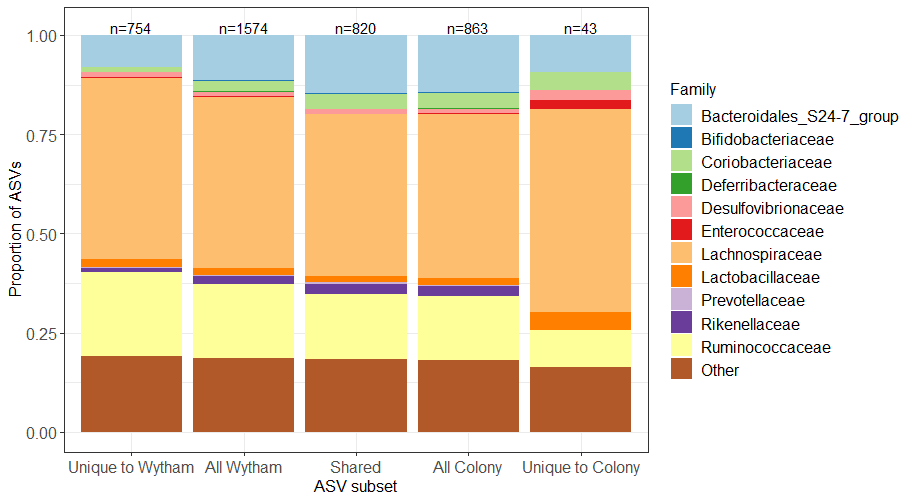

**Figure S4. Prevalence and persistence of bacteria across/within individual hosts in Wytham and Silwood.** Prevalence across individuals was calculated for each ASV using one random faecal sample per individual (n=178 in Wytham and n=75 in Silwood) and averaging the obtained value across 100 iterations. Persistence within individuals caught 3 or more times was averaged across individuals (n=57 in Wytham and n=39 in Silwood) per ASV. **a)** The relationship between prevalence and persistence of ASVs is plotted with a 1:1 dashed reference line (black). ASVs are coloured by the bacterial order they belong to. **b)** The proportion of ASVs belonging to different bacterial orders according to their position along the prevalence/persistence axis. ASVs are divided into those that were either under or over 50% prevalent and persistent as indicated by the red dashed lines in **a)** and the taxonomy of these subsets is compared to the overall taxonomic distribution.

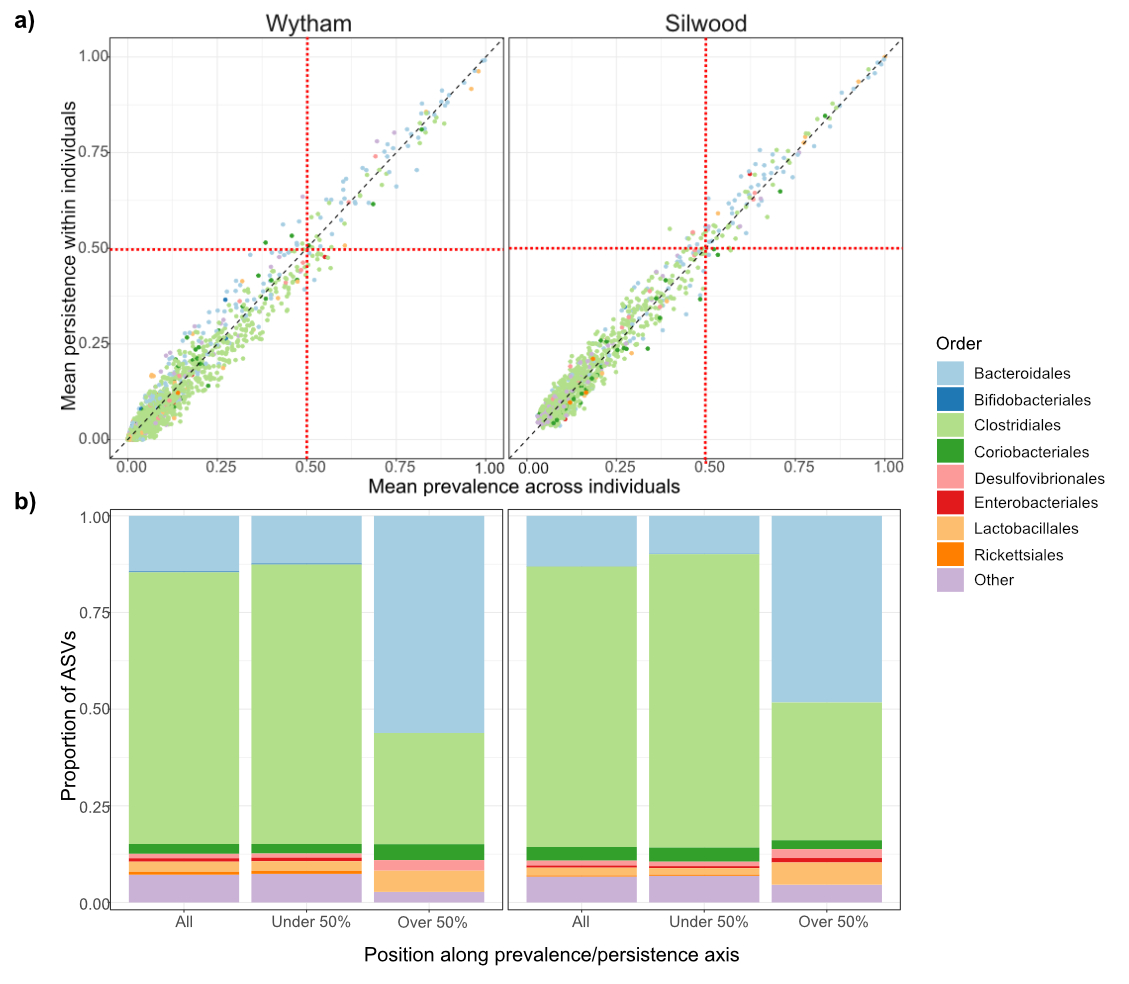

**Figure S5. Seasonal patterns in the Wytham wood mouse population age structure.** The proportion of captures per month in each year from 2015-2018 in Wytham (n=448) that were recorded as adults (A), sub-adults (SA) or juveniles (J). An increase in the proportion of young mice in the population can be seen between late Summer and Autumn.
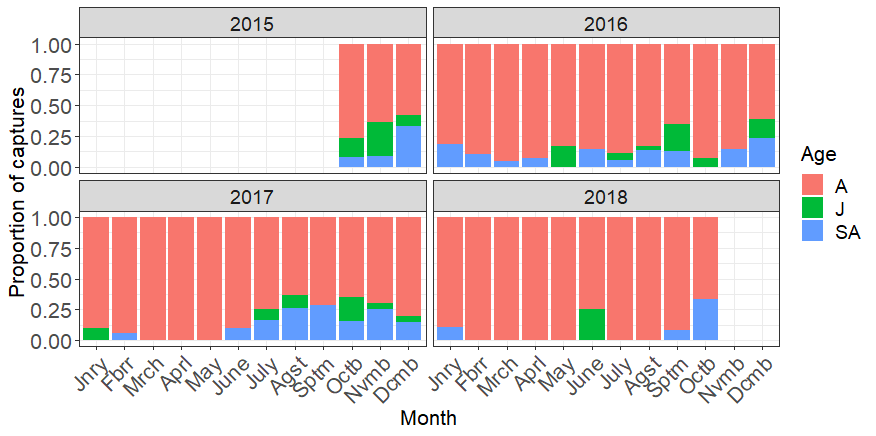

**Figure S6. Seasonal restructuring of the wood mouse gut microbiota using the Jaccard distance.** Wytham mice were sampled over 3 years between October 2015-18 (left, blue), Silwood mice were sampled for one year between November 2014-15 (right, green). Plots are derived from general additive mixed models (GAMMs) and showed predicted values and 95% confidence intervals for a smoothed day of year term within each population, controlling for repeated measures and including host-associated and methodological covariates (Wytham: n=328, approximate significance of smoothed day of year term F=520.2, edf=8.439, p<0.001; Silwood: n=185, approximate significance of smoothed day of year term F=96.3, edf=12.69, p<0.001. All host-associated covariates were non-significant (p>0.05).

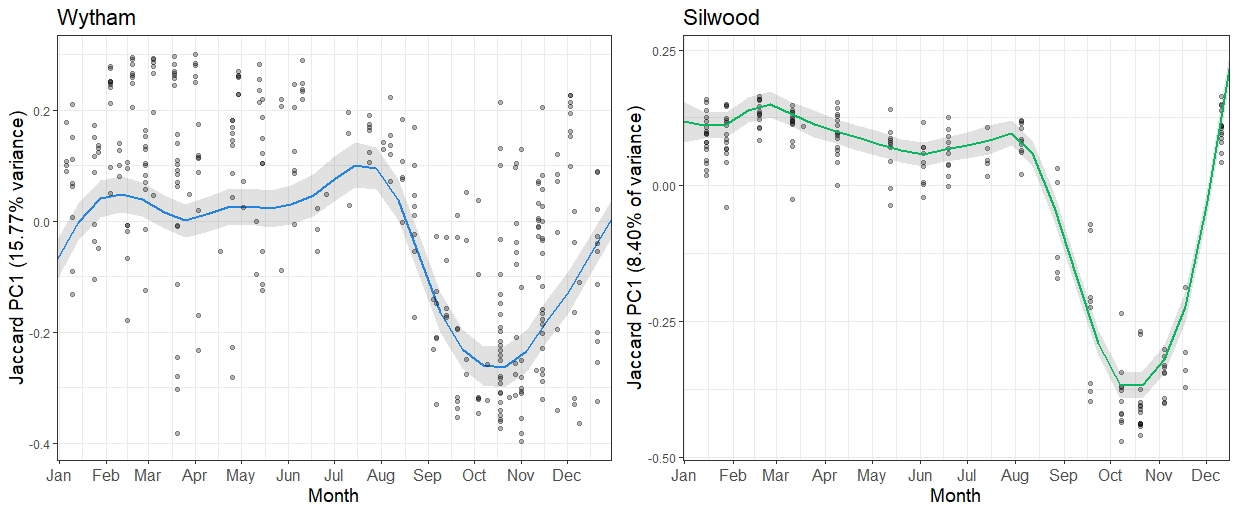

**Figure S7. Sankey plots showing the seasonal fluxes of microbial taxa in and out of the wild populations.** The presence/absence of ASVs in each season/year per population were used to visualise the flux of ASVs appearing and re-appearing between seasons from winter-autumn. The width of the bars are proportional to the total number of ASVs. Sankey plots were made using the networkD3 R package. (O) denotes the portion of ASVs that were detected in a previous season, (N) denotes ASVs that were first detected in that season.

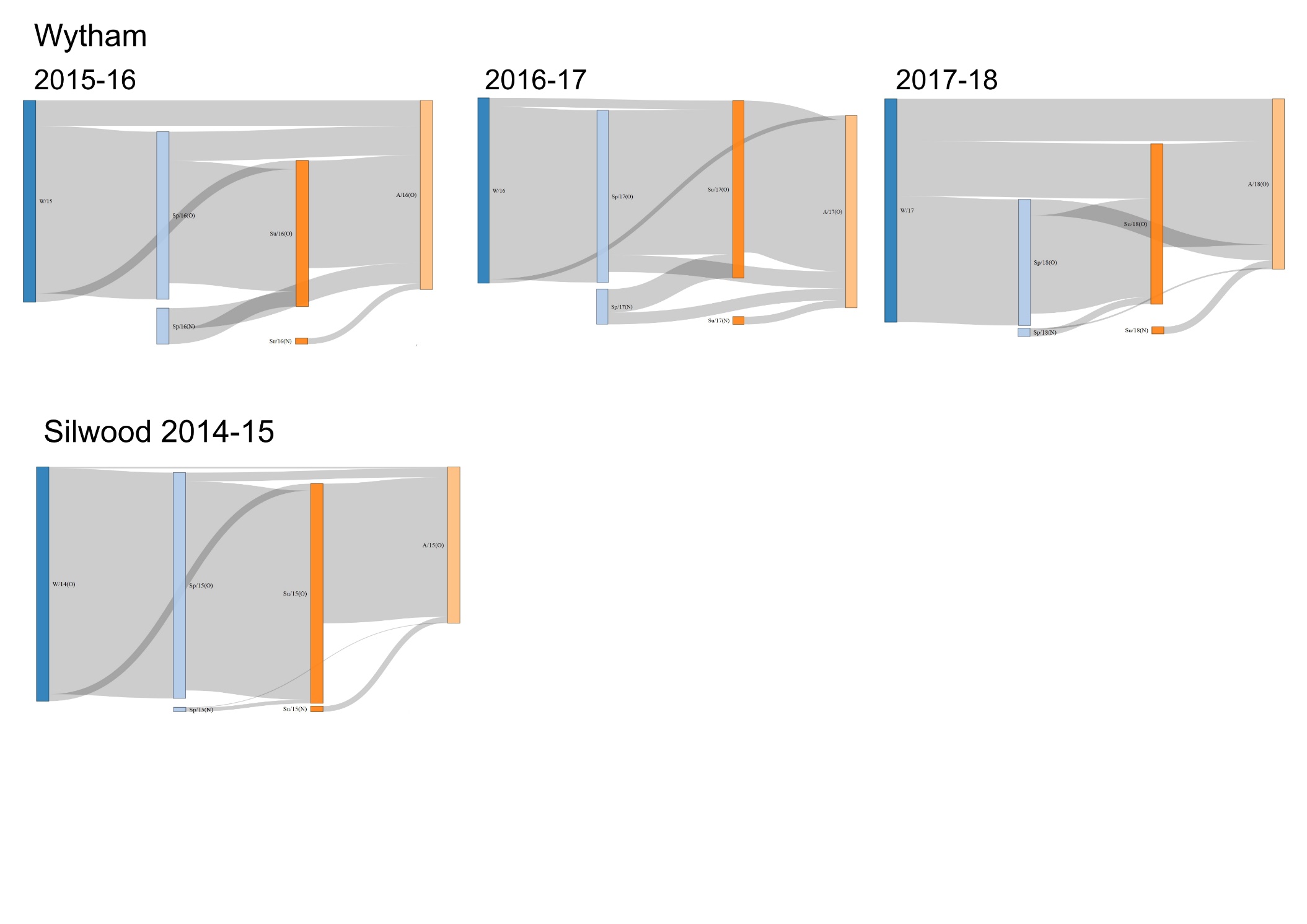

**Figure S8. Changes in family-level gut microbiota composition during seasonal restructuring.** Two wild wood mouse populations were sampled longitudinally for 3 years (Wytham, October 2015-18) and one year (Silwood; November 2014-15) and each showed consistent shifts in the first axis of a Bray-Curtis ordination (PC1) per population between July, October and February. The relative abundance of bacterial families per month are shown with the colours indicating the taxonomy at the Family level.

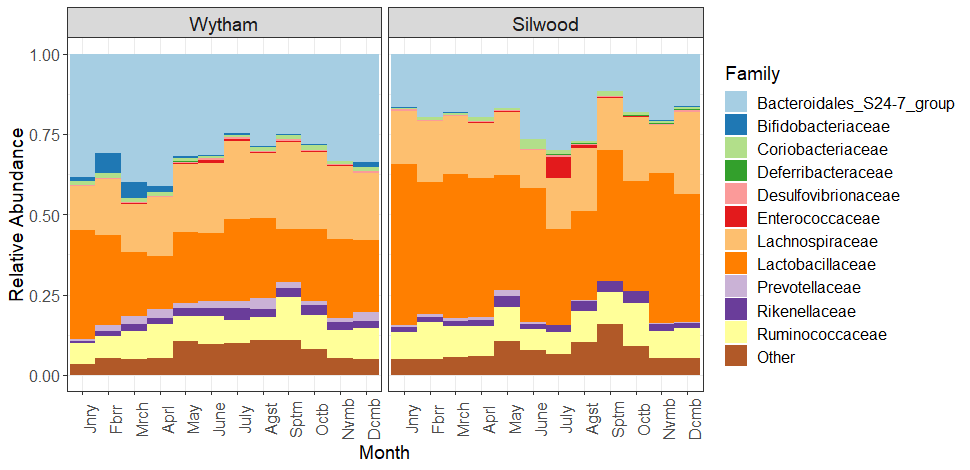

**Figure S9. Seasonality in Bray-Curtis PCoA axes.** The first three axes (PC1-3) of a Bray-Curtis PCoA of gut microbiota structure in two separate wood mouse populations sampled longitudinally for 3 years (Wytham, n=448) and one year (Silwood, n=253) are plotted against day of the year to explore seasonal patterns. The percentage variation in gut microbial community structure is shown in brackets next to the name of each axis. The temporal patterns were visualised using loess smoothing regression lines.

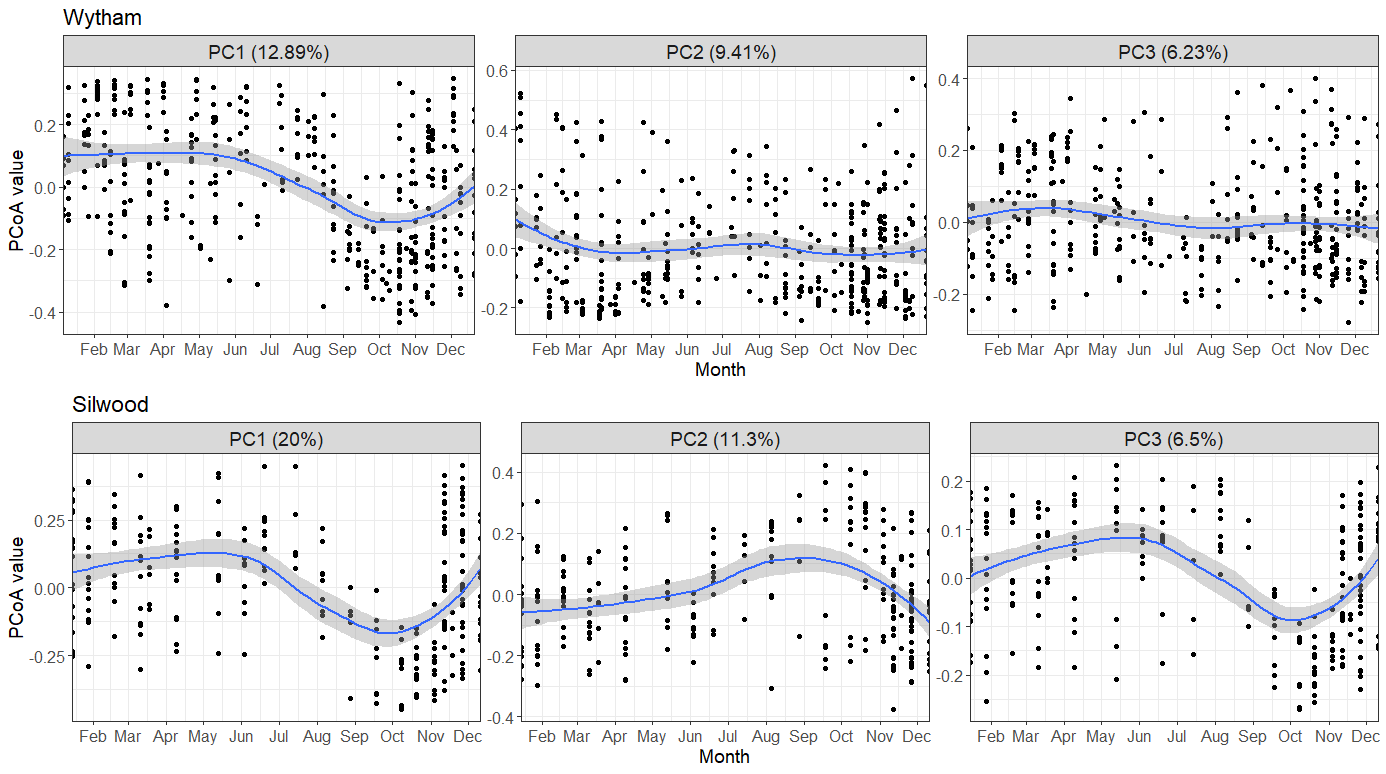

**Figure S10. Gut microbial richness is highest in early Spring compared to the rest of the year in Silwood.** Seasonal dynamics in estimated gut community richness in Silwood mice sampled for one year (November 2014-15). Predicted values and 95% confidence intervals for the smoothed day of the year from generalised additive mixed models (GAMMs) are plotted (adjusted R^2^=0.14, approximate significance of smooth term; F=1.767, edf=4.722, p<0.001, and p>0.05 for all host covariates).

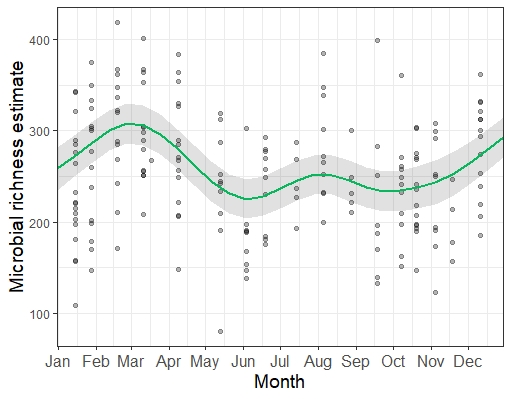

**Figure S11. Bacterial ASVs associated with the second and third PCoA axes in Silwood.** The second and third axes from a Bray-Curtis PCoA on samples collected from the Silwood mouse population (n=253) explained 11.3% and 6.5% variation in community structure, respectively, and showed seasonal patterns similar to the main PC1 in both Wytham and Silwood populations (Fig S10). Random Forest regression was used to determine the association of individual bacterial ASVs to variation along these axes. The random forest regressions could predict 86.06% variation in PC2 and 84.33% variation in PC3. ‘IncNodePurity’ was used as a measure of feature importance per PCoA axis, and the bacterial families of the top 30 ASVs are indicated by colour.

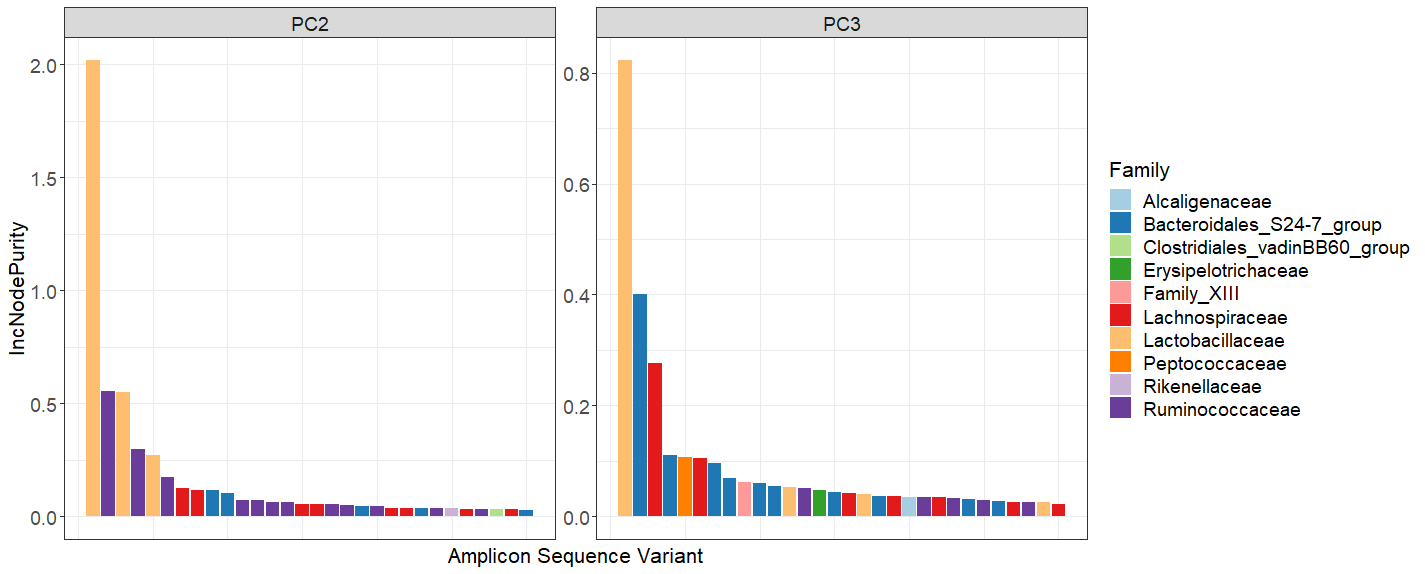

**Table S2. Results from GAMMs assessing seasonal patterns of change in gut microbiota composition.** Wytham mice were sampled over 3 years between October 2015-18 and Silwood mice were sampled for one year between November 2014-15. GAMMs were used to show the population-level seasonal trend in PC1, the first axis of a principal coordinates analysis on Bray-Curtis dissimilarity. s(Date) indicates day of the year as a smoothed term, and for this term degrees of freedom are estimated (edf) and p-values approximate. R^2^  values indicate the proportion of variance explained by the smoothed date term in each model. Models included individual ID as a random factor (which explained 26.75% and 1.79% variation in Wytham and Silwood respectively), as well as host-associated and methodological fixed effects. Terms significant in either model are shown in bold.

|  | **Wytham (n=328)** | | | | **Silwood (n=185)** | | | |
| --- | --- | --- | --- | --- | --- | --- | --- | --- |
| **Variable** | df | F | p | R^2^ | df | F | p | R^2^ |
| **s(Date)** | 9.98 | 6.35 | **<0.001** | 0.492 | 6.80 | 16.47 | **<0.001** | 0.479 |
| **Year** | 3 | 13.203 | **<0.001** |  |  |  |  |  |
| Sex | 1 | 0.745 | 0.389 |  | 1 | 0.814 | 0.368 |  |
| Reproductive status | 1 | 0.152 | 0.697 |  | 1 | 0.781 | 0.378 |  |
| Age | 2 | 1.206 | 0.301 |  | 2 | 1.516 | 0.223 |  |
| Body mass (g) | 1 | 1.888 | 0.170 |  | 1 | 0.537 | 0.465 |  |
| Body condition | 4 | 0.402 | 0.807 |  | 5 | 1.031 | 0.401 |  |
| MiSeq run | 3 | 1.920 | 0.126 |  |  |  |  |  |

**Table S3. Examining the association between host and environmental factors with gut microbiota variation using partial RDAs.** Partial redundancy analyses (pRDA) were performed on Hellinger-transformed gut microbiota community data from two wild populations, Wytham (n=328) and Silwood (n=241) using temporal, methodological and host terms as constraints. Individual ID was included as a condition to control for repeated measures. Permutation test results for the marginal effects of the constraining terms under the reduced model are shown. Overall model significance for Wytham; F=1.588_(27,160)_, p=0.001, adjusted R^2^=0.061, Silwood; F=1.579_(21,145)_, p=0.001, adjusted R^2^=0.051. In Silwood, the host sex was collinear (redundant) with other constraining terms and therefore results from the permutation tests do not include sex in this population.

|  | **Wytham** | | | | **Silwood** | | | |
| --- | --- | --- | --- | --- | --- | --- | --- | --- |
| **Variable** | df | F | p | Variance | df | F | p | Variance |
| **Read Count** | 1 | 1.389 | 0.063 | 0.002 | 1 | 1.087 | 0.271 | 0.002 |
| **MiSeq run** | 3 | 1.311 | **0.039** | 0.006 |  |  |  |  |
| **Month** | 11 | 1.496 | **0.001** | 0.023 | 11 | 1.730 | **0.001** | 0.029 |
| **Year** | 3 | 1.897 | **0.001** | 0.008 |  |  |  |  |
| **Sex** | 1 | 1.006 | 0.492 | 0.001 |  |  |  |  |
| **Reproductive status** | 1 | 0.819 | 0.785 | 0.001 | 1 | 1.212 | 0.157 | 0.002 |
| **Age** | 2 | 1.153 | 0.183 | 0.003 | 2 | 1.014 | 0.440 | 0.003 |
| **Body mass** | 1 | 1.146 | 0.236 | 0.002 | 1 | 1.427 | 0.052 | 0.002 |
| **Body condition** | 4 | 1.001 | 0.458 | 0.006 | 5 | 1.192 | 0.079 | 0.009 |

**Table S4. Rate of change in within-individual gut microbiota variation.** Intra-individual variation decays with increased sampling intervals in a non-linear way. Pairwise community similarity (1- Bray-Curtis dissimilarity) was calculated between pairs of samples collected from the same individual host, in mice that were captured three or more times in two separate populations (Wytham; n=277 samples from 57 hosts and Silwood; n=197 samples from 39 hosts). The rate of change was calculated as the community similarity value divided by the sampling interval (in days) per pairwise comparison. These rates of change were summarised across different time intervals; less than 1 month, 1-3 months, 3-6 months and 6-12 months. Values are mean±se of the rate of change, and N=number of within-individual pairwise comparisons used per summary.

|  | Wytham | | Silwood | |
| --- | --- | --- | --- | --- |
| Time interval | Rate of change | N | Rate of change | N |
| < 1 month | 0.0190±0.0109 | 418 | 0.0246±0.0121 | 228 |
| 1-3 months | 0.0057±0.0035 | 520 | 0.0071±0.0036 | 382 |
| 3-6 months | 0.0021±0.0011 | 292 | 0.0032±0.0012 | 238 |
| 6-12 months | 0.0008±0.0032 | 40 | 0.0018±0.0005 | 72 |
